# Milieu matters: An *in vitro* wound milieu to recapitulate key features of, and probe new insights into, polymicrobial biofilms

**DOI:** 10.1101/2021.01.07.425734

**Authors:** Snehal Kadam, Vandana Madhusoodhanan, Devyani Bhide, Rutuja Ugale, Utkarsha Tikhole, Karishma S Kaushik

**Author notes:** these authors contributed equally (Rutuja Ugale, Utkarsha Tikhole).

## Abstract

Bacterial biofilms are a major cause of delayed wound healing. Consequently, the study of wound biofilms, particularly in host-relevant conditions, has gained importance. Most *in vitro* biofilm studies employ refined laboratory media to study biofilms, conditions that are not relevant to the infection state. To mimic the wound milieu, *in vitro* biofilm studies often incorporate serum or plasma in growth conditions, or employ clot or matrix-based biofilm models. While incorporating serum or plasma alone is a minimalistic approach, the more complex *in vitro* wound models are technically demanding, and poorly compatible with standard biofilm assays. Based on previous reports of clinical wound fluid composition, we have developed an *in vitro* wound milieu (IVWM) that includes, in addition to serum (to recapitulate wound fluid), matrix elements and biochemical factors. In comparison with Luria-Bertani broth and Fetal Bovine Serum (FBS), the IVWM was used to study planktonic growth and biofilm features, including interspecies interactions, of common wound pathogens, *Staphylococcus aureus* and *Pseudomonas aeruginosa*. We demonstrate that the IVWM recapitulates widely reported *in vivo* biofilm features such as metabolic activity, increased antibiotic tolerance, 3D structure, and interspecies interactions for single- and co-species biofilms. Further, the IVWM is simple to formulate, uses laboratory-grade components, and is compatible with standard biofilm assays. Given this, it holds potential as a tractable approach to study wound biofilms under host-relevant conditions.

## Introduction

Wound healing is mediated by several host factors, including inflammatory, immune and biochemical components [1,2]. Following injury, a protein-rich fluid leaks into the wound, which along with cellular and matrix elements, results in a characteristic wound milieu [3,4]. Interplay across various factors is reflected in this milieu, which is known to influence progression and outcome of the wound state [5–8]. Microbial infections are the single-most important cause of delayed wound healing [2,9]. In the wound bed, bacteria form biofilms, which are polymicrobial communities enmeshed in a self-produced extracellular matrix [10–12]. Biofilms in wounds are typically observed as microscopic bacterial aggregates on the surface of, or embedded in, host tissue [13–17], and in the presence of the wound milieu.

Hitherto, the study of biofilms in wounds has typically relied on *in vivo* animal systems or *in vitro* laboratory studies [18,19]. *In vivo* systems are limited by the fact that they are not widely-available, and pose technical and ethical challenges. On the other hand, the majority of *in vitro* biofilm studies employ laboratory media [20,21] (such as refined protein broths) to grow biofilms, and analyze effects of antimicrobial treatments. However, the composition of laboratory media is not relevant in the context of the wound infection state. Recognizing this, recent studies have incorporated serum or plasma in *in vitro* growth conditions, to more closely represent the host milieu [22–27]. This is relevant given that wound fluid has been shown to resemble the biochemical and nutrient profile of serum [22]. However, the wound milieu is more complex, and includes additional host factors and matrix elements [28]. To recapitulate this, clot and matrix-based *in vitro* wound biofilm models have been developed that more closely mimic *in vivo* conditions [22,27,29–31]. However, these models are technically demanding, low-throughput, and poorly compatible with standard biofilm assays.

In this study, we have developed a simple *in vitro* wound milieu (IVWM) that includes, in addition to serum (to recapitulate wound fluid), matrix elements such as collagen, fibrinogen and fibronectin, and host factors such as lactoferrin and lactic acid. The formulation of the milieu is based on the composition of clinical wound fluid, as reported across previous studies [32,33]. We employ this composite milieu to study planktonic growth, biofilm features, and interspecies interactions of common wound pathogens, *Staphylococcus aureus* and *Pseudomonas aeruginosa* [34]. Using laboratory media (Luria-Bertani broth) and fetal bovine serum (FBS) for comparison, we demonstrate that the *in vitro* wound milieu recapitulates key *in vivo* biofilm features such as biomass formation, metabolic activity, antibiotic tolerance, three-dimensional structure, and interspecies interactions. While expectedly different from laboratory media, we find that these features are distinct from that observed with serum alone. Notably, the impact of the IVWM on co-species growth of the pathogens differs from that in serum, and similar to *in vivo* conditions appears to provide an advantage to *P. aeruginosa* [18,19]. Further, the IVWM is easy to formulate, high-throughput, and is compatible with standard biofilm assays.

## Materials and Methods

### Bacterial strains and growth conditions

All experiments were carried out using fluorescently tagged strains of *Pseudomonas aeruginosa* (PAO1-pUCP18, mCherry [35]) and *Staphylococcus aureus* (Strain AH 133-pAH13, GFP [36]). These strains were a gift from Dr. Kendra Rumbaugh (Texas Tech University Health Science Center, Lubbock, TX). Selection for SA-GFP was done with 10 μg/ml erythromycin and for PAO1-mCherry with 100 μg/ml ampicillin on Luria-Bertani (B) agar plates and in overnight LB broth cultures. Strains were streaked onto LB agar (Sigma) and incubated overnight at 37°C. Isolated colonies were grown in Luria-Bertani (LB) broth overnight under shaking conditions at 37°C, unless otherwise stated.

### Preparation of *in vitro* wound milieu

An *in vitro* host milieu (IVWM) was prepared with sterile fetal bovine serum (FBS) (Thermofisher Scientific) as the base component. Other components added included sterile rat tail collagen (Sigma, 50 μg/ml), lactoferrin (Sigma, 2 mg/mL stock prepared by dissolving in 1X PBS (pH 7.4) and filter sterilized), fibronectin (Sigma, 1 mg/ml stock solution prepared using autoclaved distilled water), fibrinogen (Sigma, 0.9% NaCl (prewarmed at 37°C) was used to prepare a stock solution of 10 mg/ml and filter sterilized) and lactic acid (Sigma, 11.4 M stock concentration). Collagen and lactoferrin were stored at 4°C. FBS, fibronectin and fibrinogen were stored at −20°C. Lactic acid was stored at room temperature. Components were either purchased sterile or filter sterilized using a 0.22 μm syringe filter. The components were combined in concentrations given in Table 1 to result in the final IVWM. The IVWM was freshly prepared each time, and was used immediately after use (not stored). The pH of the IVWM was measured using a pH probe and pH strip. Specific gravity of IVWM was calculated as the relative weight of IVWM compared to the weight of an equal volume of distilled water [37].

**Table 1:**
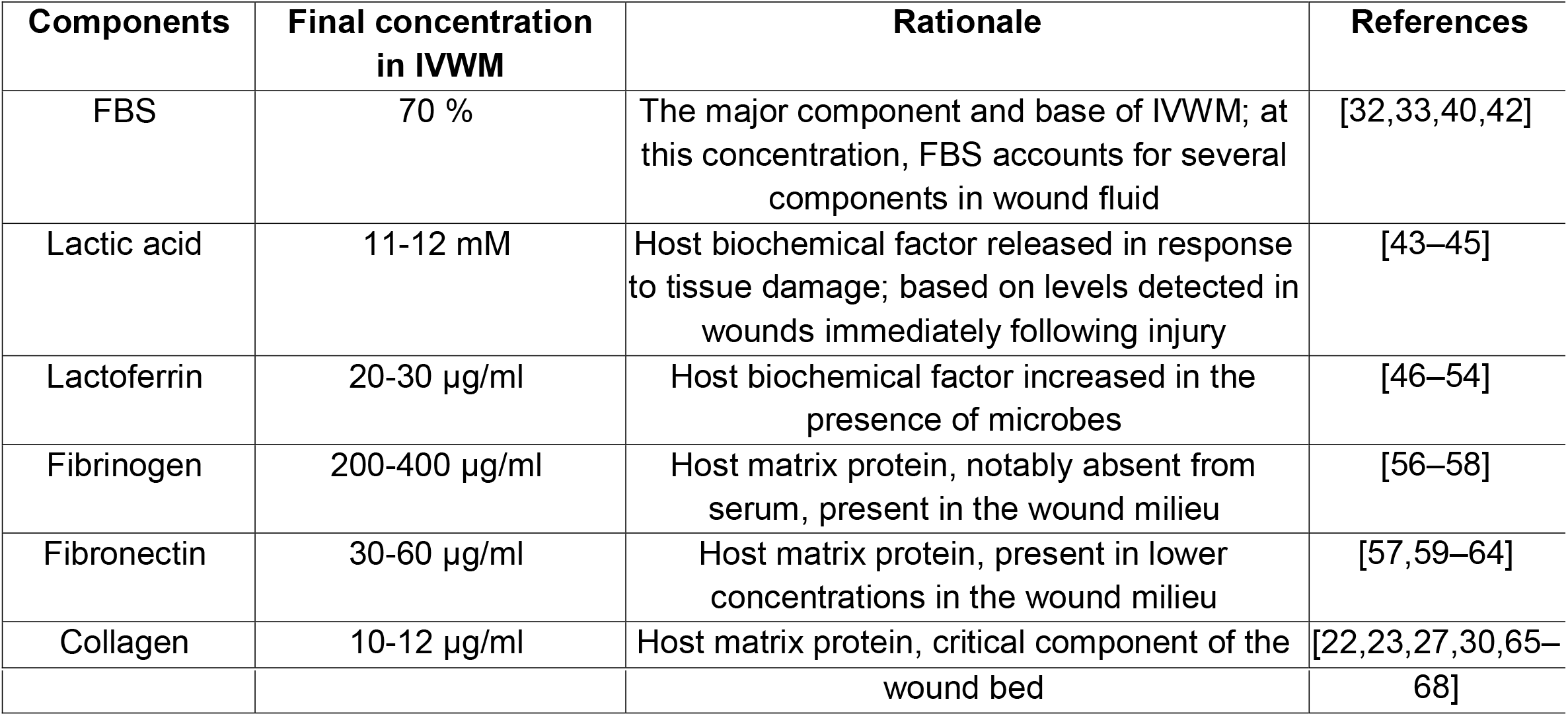
Composition of the *in vitro* wound milieu (IVWM) and rationale for inclusion of components

### Planktonic growth

#### For growth curves in LB

Overnight cultures of *S. aureus* and *P. aeruginosa* were grown in LB broth under shaking conditions at 37°C. The next day, cultures were quantified by measuring optical density (O.D.) at 600 nm with a multimode microplate reader (Tecan Infinite 200 PRO). To set up growth curves, overnight cultures were diluted in sterile LB broth (1:100) and 100 μL of the diluted culture (consisting of ~10^5^ cells) was added per well, in replicates of three, to a sterile, transparent, round bottom, untreated 96-well polystyrene plate. Uninoculated LB was used as a control. Plates were incubated in the multimode microplate reader (Tecan Infinite 200 PRO) at 37°C with shaking in orbital mode (2 mm amplitude), and O.D. was measured at 600 nm every 30 minutes for 12-14 hours.

#### For growth curves in FBS and IVWM

Growth curves in FBS and IVWM were done using overnight cultures set up in FBS. Briefly, each bacterial colony was inoculated in FBS and incubated overnight at 37°C under shaking conditions. The next day, cultures were quantified by measuring O.D. at 600 nm with a multimode microplate reader (Tecan Infinite 200 PRO). To set up growth curves, each culture was diluted in sterile FBS or IVWM (1:100) and 100 μL of this diluted culture (consisting of ~10^5^ cells) was added per well, in replicates of three, to a sterile, transparent, round bottom, untreated 96-well polystyrene plate. Uninoculated FBS or IVWM was used as a control. The plate reader was set to a temperature of 37°C with shaking in orbital mode (2 mm amplitude), and absorbance was measured at 600 nm every 30 minutes for 12-14 hours.

For all growth curves, O.D. versus time was plotted and growth rates (doubling times) were calculated.

### Colony Forming Units (CFUs)

In order to quantify the proportion of living cells in the planktonic cultures, alone and under co-species conditions, colony count assays were carried out. Overnight cultures in LB, FBS and IVWM were diluted to ~10^6^ cells/ml in respective media. From these diluted cultures, 100 μL (containing ~10^5^ cells) was added to 100 μL of the respective media in a fresh tube (for single-species cultures). For co-species cultures, 100 μL each of the diluted culture of *S. aureus* and *P. aeruginosa* (~10^5^ cells of each strain) were added together in a single fresh tube. All conditions were set up in at least 3 replicates. Cultures were incubated under shaking conditions at 37°C for 7 hours. Cultures were then diluted in LB to different dilutions and plated on selective media *Pseudomonas* isolation agar BioVeg (SRL Chemicals) and *Staphylococcus* Medium 110 (SRL Chemicals). Plates were incubated overnight for 24 hours at 37°C. Based on protocols using *Staphylococcus* Medium 110, these plates required incubation for upto 36-48 hours to obtain visible countable colonies.

### Biofilm formation

Overnight cultures *S. aureus* and *P. aeruginosa* were each diluted to ~10^6^ cells/ml in LB, FBS and IVWM. From these diluted cultures, 50 μL (~10^5^ cells) was added in multiple replicates (at least three) to a transparent, round bottom 96-well polystyrene plate, unless otherwise stated. To these wells, 50 μL of the media in which biofilm formation was to be tested was added, to maintain a constant volume of 100 μL.

For polymicrobial biofilms, individual cultures of *P. aeruginosa* and *S. aureus* were diluted to ~10^6^cells/ml (as above). From these diluted cultures, 50 μL (~10^5^ cells) of each was added to wells in at least three replicates to make a total volume of 100 μL. Biofilms were allowed to grow under static conditions at 37°C for 24 hours. These preformed biofilms were used to measure metabolic activity (XTT assay) and to visualize the biofilm (confocal microscopy).

### XTT assay for biofilm metabolic activity

Pre-formed biofilms of *S. aureus* and *P. aeruginosa* grown for 24 hours under different conditions (as described above) were washed once with LB after removing suspended media. Menadione solution (7 mg/ml) was diluted 1:100 in sterile distilled water. A mixture of LB:XTT:Menadione in 79:20:1 ratio was freshly prepared [38], and 150 μL of this was added to each well. The plates were covered in aluminum foil and incubated for 4 hours at 37°C (static). From each well, 100 μl was transferred to a new 96-well plate and absorbance was measured at 492 nm. In order to evaluate the presence or absence of any interactions in co-species biofilms, the co-species biofilm absorbance value was compared to an ‘expected’ value (calculated as the additive absorbance value of the single-species biofilms).

### Antibiotic Susceptibility of Pre-formed Biofilms

Pre-formed biofilms of *S. aureus* and *P. aeruginosa* were grown for 24 hours as previously described. Briefly, overnight cultures grown in LB were diluted in the respective media (LB, FBS, IVWM) in which antibiotic susceptibility was to be tested. From these diluted cultures, ~10^5^ cells were added (in triplicates) to each well of a 96-well plate, and the plate was incubated for 24 hours at 37°C. After 24 hours, the suspended media was gently removed and the biofilms were washed once with LB. Antibiotics were diluted to varying concentrations (0-64 μg/mL for tobramycin and 0-512 μg/mL for vancomycin) in the respective media in which biofilm susceptibility was to be tested (LB, FBS, IVWM) and 100 μl of the antibiotic-media solution was added into the wells. Wells with untreated biofilms (no antibiotic added) were also included. Uninoculated media was used as a control. Plates were incubated at 37°C for 24 hours. After 24 hours, the XTT assay was performed to quantify viability (as described above). The concentration resulting in 80% reduction in biofilm metabolic activity (representing living cells), compared to the untreated biofilms, was considered as the MBEC_80_ value for that particular antibiotic.

### Biofilm Visualization using Confocal Microscopy

To visualize *in situ* three-dimensional biofilm structure, 24-hour old undisturbed biofilms were set up as previously described in LB, FBS or IVWM (as single-species and co-species). Briefly, LB overnight cultures of *P. aeruginosa* and *S. aureus* were diluted in the respective media in which biofilm structure was to be observed and mixed in a 1:1 ratio for co-species biofilms. After incubation at 37°C for 24 hours, the biofilms were directly examined with confocal laser scanning microscopy (Leica SP8 Spectral CLSM). To enable this direct visualization, biofilms were grown in 96-well, black polystyrene tissue-culture treated flat-bottom plates with a transparent bottom (Corning). These plate specifications allow visualization of the biofilm structure in the well and minimum interference of fluorescence signals from neighboring wells. The tissue-culture treatment imparts an overall negative charge to the surface, and results in a hydrophilic surface, and this treatment (achieved by corona discharge) is known to reduce the attachment of negatively charged bacteria (such as *P. aeruginosa* and *S. aureus*). In doing so, this enables the study of the role of the media conditions (host components in FBS and IVWM) in biofilm formation.

Wells were imaged using a 488 nm laser for excitation and a 496–551 nm emission filter for *S. aureus*-GFP, and a 561 nm laser for excitation and a 590–625 nm emission filter for *P. aeruginosa*-mCherry. To visualize the 3D architecture of the biofilms across the entire well, an 8×8 tile scan approach was used with an overlap of 30% and a Z-stack step size of 10 μm. The images were processed and reconstructed in the Leica Application Suite (LAS) software. Mean intensity measurements were carried out using the LAS software. Biofilm thickness was measured using ImageJ.

## Results and Discussion

### Development of an *in vitro* wound milieu (IVWM) that mimics host conditions

Following injury, the wound bed is bathed in protein-rich exudate, which along with additional host elements, results in a characteristic wound milieu [1,3,4,39]. The composition of wound fluid has been widely reported to resemble that of serum [32,33,40], with several *in vitro* wound studies using serum to mimic wound conditions [22,27,30,41]. To develop an *in vitro* wound milieu (IVWM), we used fetal bovine serum (FBS) as the base component, to which relevant host matrix and biochemical factors were added. Components were chosen based on previous reports of biochemical analyses across clinical wound fluids [32,33], as well as their identified roles in the wound bed.

Based on analysis of wound fluid composition [32], we decided to use 70% FBS as the base component of the IVWM, since at this concentration the levels of multiple components in serum fit into the range of values for that component in wound fluid [32,42]. While the concentrations of several biochemical factors in serum and wound fluid are similar [32], the wound milieu is also characterized by the presence of additional host-derived biochemical factors. In the initial inflammatory state of wound repair, the wound microenvironment is characterized by increased levels of lactate and lactoferrin. Lactate in the wound microenvironment is a result of local tissue damage, and increased concentrations are reflected in the local milieu [43–45]. While the levels of lactoferrin in the plasma and serum of healthy individuals is typically low [46–49], in the presence of microbial infections [50], increased levels are reported in the wound milieu. In addition to possessing antimicrobial properties [51–53], lactoferrin has also been shown to be important for wound re-epithelialization [54]. Given this, lactate and lactoferrin were included in the IVWM at concentrations that mimicked host conditions (Table 1).

The IVWM composition also included relevant matrix components such as fibrinogen, fibronectin and collagen [55]. While fibrinogen is present in wound tissue [56,57] and plasma [58], it is notably absent from serum. On the other hand, the matrix protein fibronectin is typically present at high concentrations in serum or plasma [59], but undergoes degradation under inflammatory wound conditions [60–62]. To mimic this, fibronectin was added to the IVWM, to resemble lower concentrations as relevant to the wound milieu [57,62–64]. Given its critical role as an extracellular matrix protein in the wound bed [22,23,27,30,65–68], collagen was also included in the IVWM (Table 1). At the concentration of collagen added, the IVWM was in liquid form (not a gel) resembling the wound fluid milieu.

The pH of the formulated IVWM was determined to be 5.25 and specific gravity was 0.966, which is similar to that measured in wound fluid [33,69,70].

### Planktonic growth of *P. aeruginosa* and *S. aureus* in laboratory media, fetal bovine serum and the *in vitro* wound milieu (IVWM)

To determine the effects of the IVWM on the planktonic growth of *P. aeruginosa* and *S. aureus* (single-species), we performed growth curves in the IVWM, and in LB medium and FBS for comparison (Figure 1). Luria-Bertani (LB) broth is widely-used to study biofilms under *in vitro* conditions [20,38,71–73], however, its composition (refined yeast extract and tryptone) poorly mimics the infection state. To recapitulate factors relevant to the wound milieu, several biofilm models incorporate fetal bovine serum (FBS), in varying concentrations, into growth conditions [22,26,74,75]. Given that concentrations of FBS employed across different studies range widely, we chose to use the maximum possible FBS concentration (100%) for comparison.

**Figure 1:**
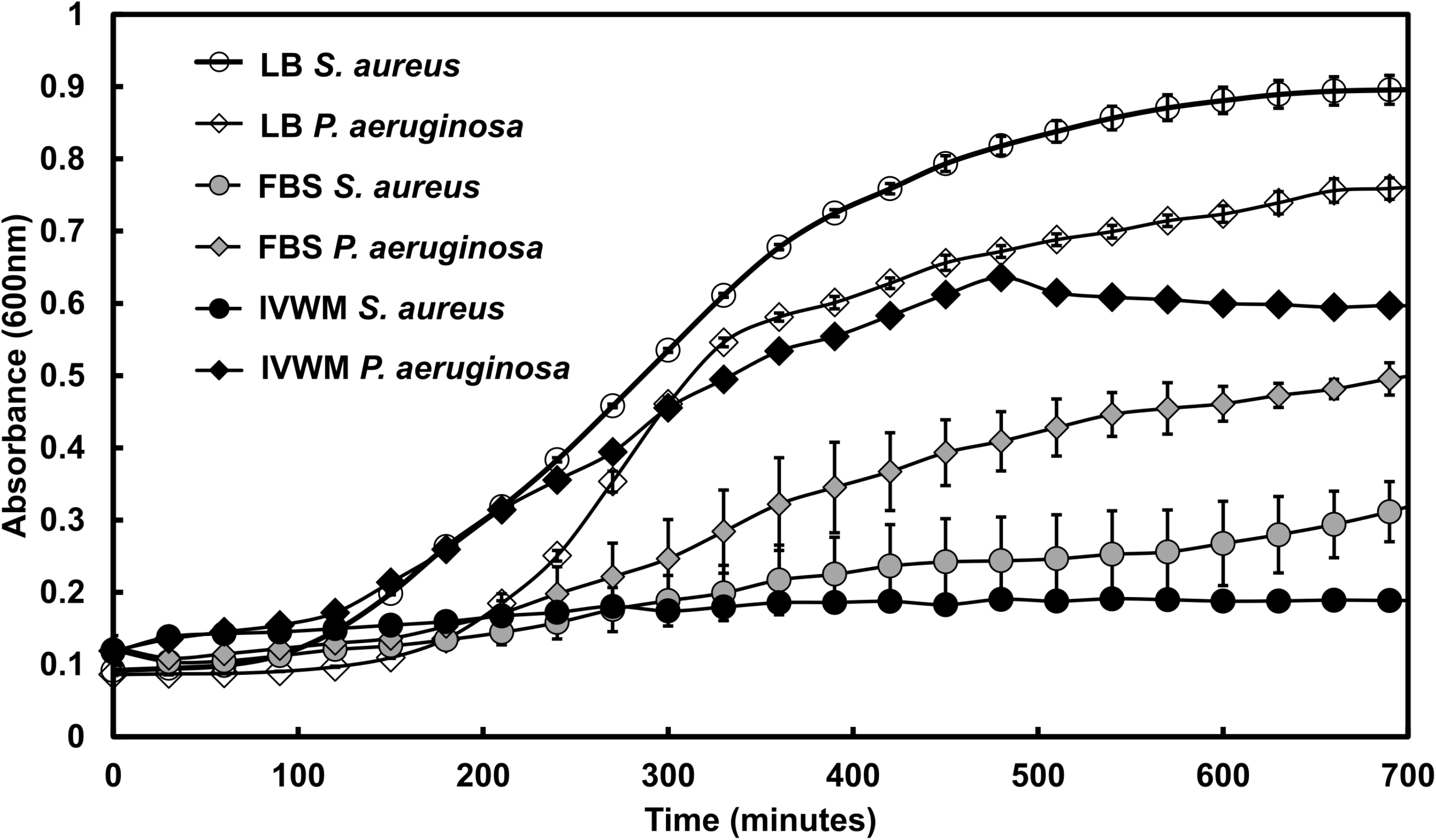
IVWM supports the planktonic growth of *P. aeruginosa*, but not of *S. aureus*. Planktonic growth curves of *Pseudomonas aeruginosa* (PAO1) and *Staphylococcus aureus* (AH133) were performed in Luria-Bertani broth (LB), Fetal Bovine Serum (FBS) and the *in vitro* host milieu (IVWM). Optical density (OD 600) was measured at intervals of 30 minutes for 12 hours. Error bars represent SEM, n=3.

When examined in LB medium, *P. aeruginosa* and *S. aureus* showed typical growth curves, with doubling times of 30±1 minutes and 28±1 minutes respectively, in accordance with previous reports [76,77]. In 100% FBS, *P. aeruginosa* was observed to grow slower as compared to growth in LB medium, with a doubling time of 56±7 minutes. On the other hand, *S. aureus* showed markedly impaired growth in FBS. In the IVWM, consisting of 70% FBS with additional matrix and host components, *P. aeruginosa* was observed to double every 43±7 minutes, which is faster than that seen in FBS alone, and cultures were also observed to enter exponential phase earlier. Similar to that observed in FBS, *S. aureus* demonstrated significantly impaired growth in the IVWM.

As compared with LB media, we find that the planktonic growth of *P. aeruginosa* and *S. aureus* are notably different under growth conditions that incorporate host factors. In the presence of FBS, both pathogens displayed markedly slower growth; components in serum are known to impair the growth of *S. aureus* [78–80]. However, when grown in IVWM, containing serum at a concentration that recapitulates wound fluid (70% FBS), and with additional matrix and biochemical factors, a greater difference in growth across the two species was observed. The IVWM was observed to better support the growth of *P. aeruginosa* (as compared with FBS alone), while *S. aureus* showed significantly impaired growth. This indicates that in the IVWM, *P. aeruginosa* has a distinct growth advantage in planktonic state, as compared with its co-pathogen *S. aureus*.

### Interspecies interactions between planktonic *P. aeruginosa* and *S. aureus* under different conditions

We next wanted to understand the effects of the IVWM, in comparison with LB and FBS, on interspecies interactions between planktonic *P. aeruginosa* and *S. aureus*. For this, *P. aeruginosa* and *S. aureus* were co-cultured under planktonic conditions in LB, FBS and IVWM (at a starting ratio of 1:1, with ~10^5^ CFU of each strain), and after 7 hours, were plated on selective media to obtain viable counts of each bacterial species. In LB media, both *S. aureus* and *P. aeruginosa* showed a decrease in viable cells in co-species conditions (Figures 2A and B), the number of viable cells recovered for *S. aureus* was ~3% and ~50% for *P. aeruginosa*, compared to when grown alone. Notably, this inhibitory effect was more predominantly observed for *S. aureus* (Figure 2B), which is in accordance to previously published reports of *P. aeruginosa* outcompeting *S. aureus* in LB medium [18,20,21,31,81,82].

**Figure 2:**
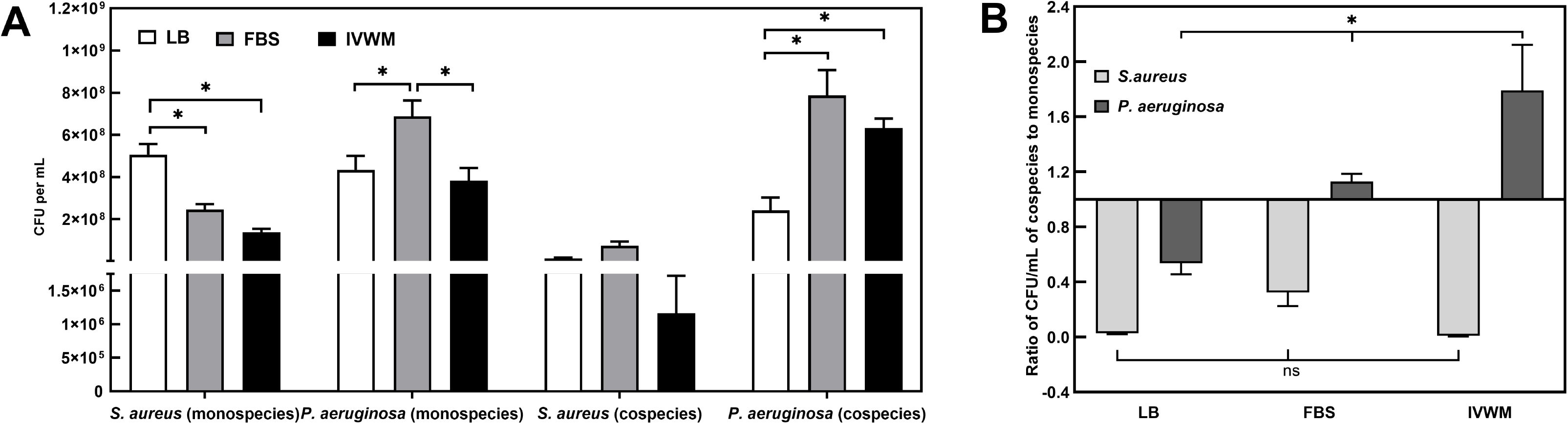
In the IVWM, interspecies interactions between planktonic *P. aeruginosa* and *S. aureus* result in *P. aeruginosa* significantly outcompeting *S. aureus*. Viable counts of planktonic *P. aeruginosa* and *S. aureus* grown under co-species conditions in Luria-Bertani broth (LB), Fetal Bovine Serum (FBS) and the *in vitro* wound milieu (IVWM) were quantified using the Colony Forming Units (CFU) technique. (A) CFUs of planktonic single-species and co-species cultures of *Pseudomonas aeruginosa* and *Staphylococcus aureus* in LB, FBS and IVWM (B) Ratio of CFU/mL of the colony counts in co-species to that in single-species planktonic cultures of *Pseudomonas aeruginosa* and *Staphylococcus aureus* in LB, FBS and IVWM. Error bars represent SEM, n=4.

In FBS and IVWM, *S. aureus* demonstrated the same trend, with reduced recovery of viable cells under co-species conditions (Figure 2). Notably, in the IVWM, viable *S. aureus* cells recovered from planktonic co-cultures were very few, representing less than 1% of the viable *S. aureus* cells in single-species cultures. However, under these conditions, the recovery of viable *P. aeruginosa* was similar or better under co-species conditions compared to that when grown alone. This effect was more pronounced in the IVWM (Figure 2A and B), with a 46-112% increased recovery under co-species conditions. It is important to note that even when grown alone, the IVWM is observed to better support the growth of *P. aeruginosa* as compared to *S. aureus* (Figures 1 and 2A). Further, in co-species conditions, interspecies interactions could result in *P. aeruginosa* inhibiting the growth of *S. aureus*, resulting in a net effect where *P. aeruginosa* significantly outcompetes *S. aureus*. This overall effect is similar to that seen under *in vivo* conditions, where in spite of co-existence between the two common wound pathogens, *P. aeruginosa* is observed to outcompete *S. aureus* in wound infections [18,31,81,83–85].

### Biofilm formation, metabolic activity and interspecies interactions of *P. aeruginosa* and *S. aureus* biofilms under different conditions

To understand the effects of the IVWM on biofilm formation, metabolic activity and interspecies interactions of *P. aeruginosa* and *S. aureus* biofilms, 24-hour biofilms (alone and as co-species) were grown in microtiter plates in IVWM, and in LB and FBS for comparison. Biofilms were then examined with the XTT assay (Figure 3), which measures bacterial metabolic activity, and is therefore a proxy for living cells in the biofilm.

**Figure 3:**
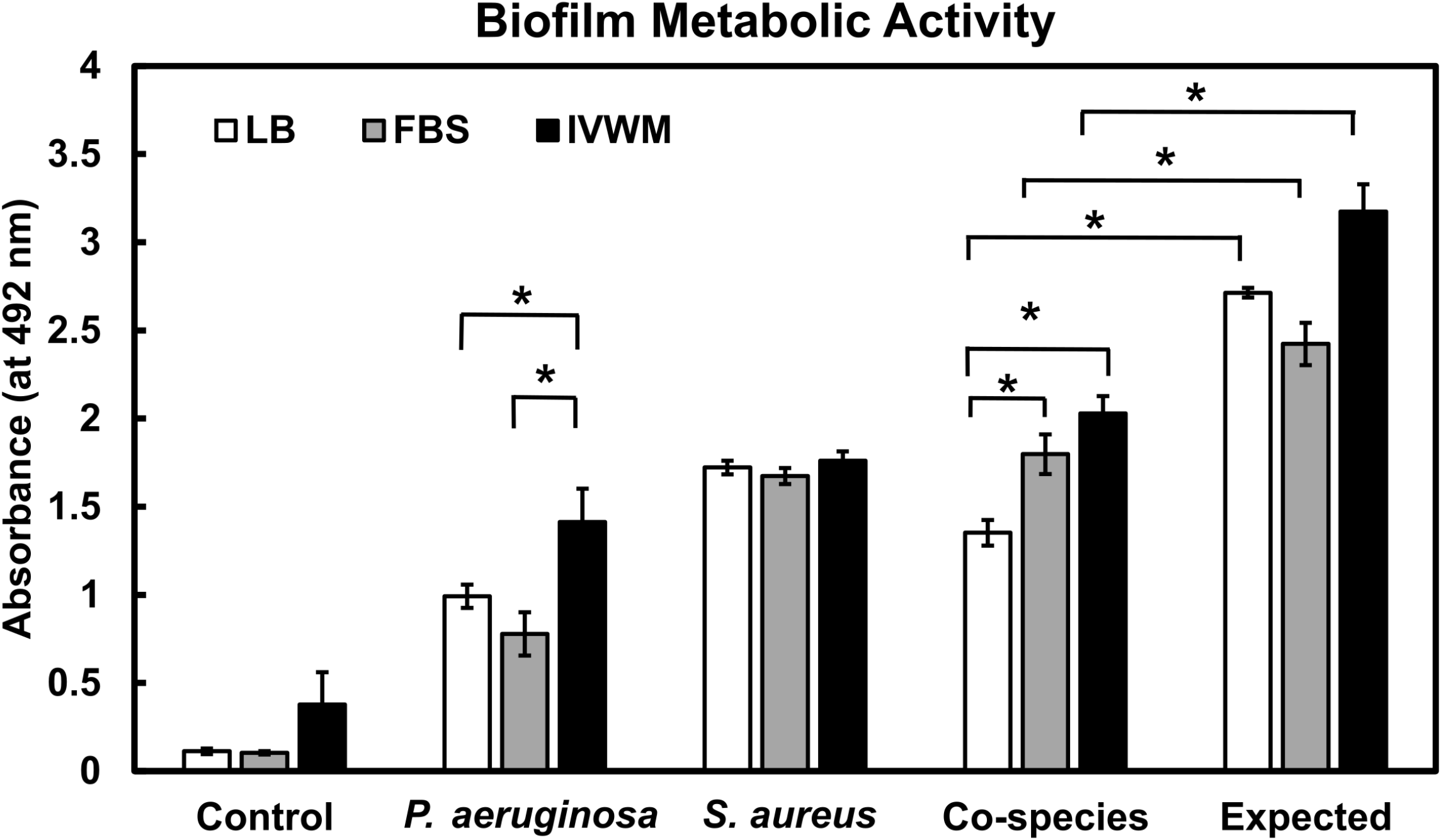
IVWM supports the formation of metabolically-active biofilms of *P. aeruginosa* and *S. aureus*, and indicates the presence of interspecies interactions. Pre-formed 24-hour biofilms of *Pseudomonas aeruginosa* and *Staphylococcus aureus* were quantified for metabolic activity, in single-species and co-species state, in LB, FBS and IVWM, by the XTT assay. Error bars represent SEM, n=3.

As previously reported across several studies [20,71], after 24 hours of growth in LB medium, *P. aeruginosa* and *S. aureus* formed biofilms that displayed the presence of metabolic activity (Figure 3). This is also observed in the presence of host components, as in FBS and IVWM, where both *P. aeruginosa* and *S. aureus* formed metabolically-active biofilms (Figure 2). Several host components, including serum, plasma, and matrix factors such as collagen, fibrinogen, fibronectin, have been shown to support the formation of *P. aeruginosa* [86–88] and *S. aureus* [89–97] biofilms. It is important to note that in our study, though *S. aureus* biofilms in FBS showed the presence of metabolic activity, when examined visually they were seen as a thin layer on the bottom and sides of the microtiter well (data not shown). This could possibly be the reason that certain previous studies using biomass staining protocols (and not metabolic activity) [23,80,96], report reduced or absent biofilm formation in the presence of FBS or plasma.

When *P. aeruginosa and* S. *aureus* biofilms were grown together in LB medium, the metabolic activity of the co-species biofilms is significantly reduced as compared with that expected (the additive value of single-species biofilms of both pathogens) (Figure 3), with the observed values being 50±12% of the expected value. Interspecies interactions between *P. aeruginosa* and *S. aureus* in biofilms are widely reported [18,20,21,34,98–100], and are influenced by several factors, including host matrix and chemical factors. In the presence of host components, as in FBS and the IVWM, the metabolic activity of co-species biofilms was similarly less than that expected (Figure 3), observed as 74±15% and 64±15% of the expected value respectively. This indicates that, even in the presence of host components, co-species biofilms of *P. aeruginosa* and *S. aureus* demonstrate the possible effects of interspecies interactions.

Overall, our results indicate that the IVWM supports biofilm formation and metabolic activity of *P. aeruginosa* and *S. aureus* biofilms, and indicates the presence of interspecies biofilm interactions.

### Antibiotic susceptibility of *P. aeruginosa* and *S. aureus* biofilms in the *in vitro* wound milieu (IVWM)

To study the effects of the IVWM of the antibiotic susceptibility of *P. aeruginosa* and *S. aureus* biofilms, pre-formed biofilms were grown for 24 hours (also in LB and FBS), and exposed to varying concentrations of tobramycin and vancomycin respectively, and assayed for antibiotic susceptibility (Table 2). The Minimum Biofilm Eradication Concentration (MBEC) was considered to be the concentration resulting in 80% of biofilm eradication (MBEC_80_).

**Table 2:** Minimum Biofilm Eradication Concentration (MBEC)* for *P. aeruginosa* and *S. aureus* under different conditions. (*The concentration resulting in 80% reduction in biofilm metabolic activity (by the XTT assay) was considered as the MBEC_80_ value for that particular antibiotic)

|  | LB | FBS | IVWM |
| --- | --- | --- | --- |
| ***P. aeruginosa***<br>**(PAO1)**<br>Tobramycin | 1 µg/mL | 8 µg/mL | 8 µg/mL |
| ***S. aureus***<br>**(AH 133)**<br>Vancomycin | > 512 µg/mL | 16 µg/mL | > 512 µg/mL |

For biofilms grown in LB media, the MBEC_80_ of *P. aeruginosa* for tobramycin was similar to the previously reported value of 1 μg/mL [38]. On the other hand, the MBEC_80_ of preformed *P. aeruginosa* biofilms grown in FBS and IVWM was determined to be 8 μg/mL, an 8-fold increase compared to LB. While the presence of serum is known to increase antimicrobial tolerance [101–103], owing to the serum-binding properties of certain antibiotics, including binding of tobramycin and serum albumin [104], this increased tolerance could also be due to the inherent properties of the biofilm formed under these conditions. The similarity of MBEC_80_ in FBS and IVWM (which contains 70% FBS) could possibly indicate the dominant role of serum in influencing the antibiotic tolerance of *P. aeruginosa* biofilms.

When grown in LB media, pre-formed *S. aureus* biofilms displayed increased tolerance, even at higher concentrations of vancomycin (MBEC_80_>512 μg/mL). This is in accordance with previously reported values, across different laboratory media conditions [73,101,105,106]. Interestingly, we find that biofilms grown in FBS demonstrate increased susceptibility to vancomycin, with a 32-fold reduction in the MBEC_80_ (8 μg/mL) in FBS, as compared with LB. Visually, *S. aureus* biofilms in FBS were observed as a very thin layer on the round-bottom surface of the microtiter wells data not shown). Despite having a high metabolic activity, we speculate that inhibitory effects of serum on *S. aureus* [80], along with the formation of thin biofilms, leads to this increased susceptibility.

When grown in IVWM, 24-hour old *S. aureus* biofilms showed increased tolerance to vancomycin, with an MBEC_80_ greater than 512 μg/mL. This value is 32-fold higher than that observed for biofilms grown in FBS alone, which is important to note given that the IVWM comprises 70% FBS, along with additional matrix and host factors. Host factors and matrix components, such as plasma [90] and fibronectin [107], have been shown to reduce the susceptibility of *S. aureus* to vancomycin. On the other hand, this effect could also be due to host components such as lactoferrin, known to have antimicrobial properties against *S. aureus* [108–112]; however, its role in the presence of multiple components such as in the IVWM, or even in serum alone, is not known.

Wound biofilms are known to display increased tolerance to antimicrobial treatments, and our results show that the host and matrix factors present in the IVWM recapitulates the increased antibiotic tolerance of both *P. aeruginosa* and *S. aureus* biofilms.

### 3D Biofilm structure of *P. aeruginosa* and *S. aureus* biofilms under different conditions

In order to visualize the 3D structure of the biofilms under different conditions, undisturbed 24-hour old biofilms of *P. aeruginosa* (PAO1-mCherry) and *S. aureus* (AH133-GFP) were imaged (as single and mixed-species) using confocal microscopy via the tilescan approach. The tilescan approach uses an 8×8 grid set to scan the entire well, and thereby enables visualization of the 3D structure across the entire biofilm. To reduce the role of surface attachment [113], given that polystyrene is not a biotic surface, and better understand the role of the media conditions in biofilm structure, biofilms were grown in tissue-culture treated 96 well plates.

When grown alone in LB media, *S. aureus* formed dense biofilms after 24 hours, with an average thickness of 92±3 μm (Figure 4A). In FBS, *S. aureus* biofilms were observed to be thinner, as compared with LB, with an average thickness of 63±4 μm (Figure 4A and C). In the IVWM, *S. aureus* formed biofilms that were 36-56% thinner than that seen in FBS and LB, with an average thickness of 40±3 μm (Figure 4A). Notably, this indicates that studies in refined protein-based media, such as LB broth, significantly overestimate biofilm thickness.

**Figure 4:**
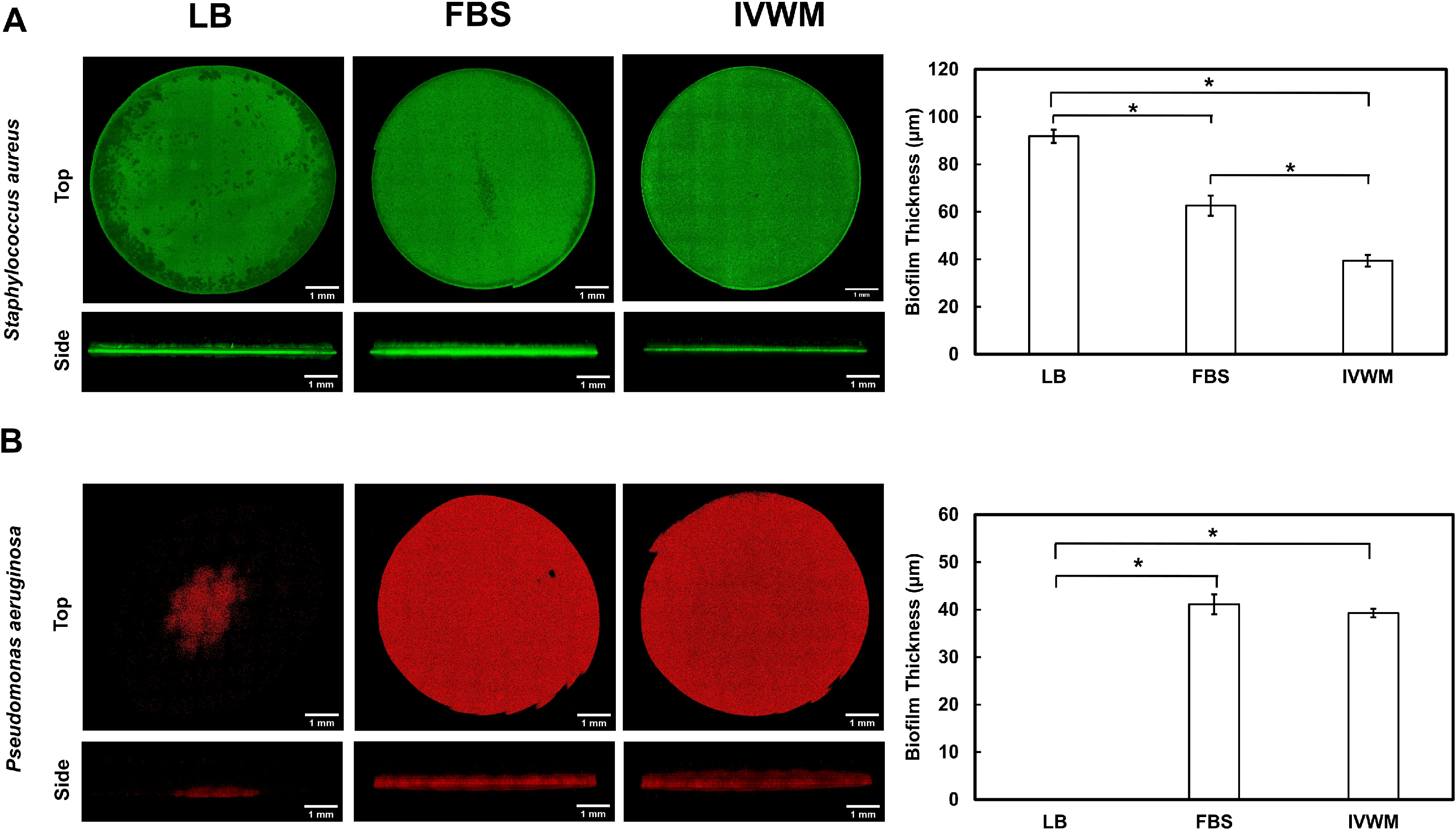
3D biofilm structure of *P. aeruginosa* and *S. aureus* single-species biofilms in IVWM are distinct from that in LB and FBS. Tile-scan confocal microscopy showing 3D structure of **(A)** *P. aeruginosa* (PAO1-mCherry) and **(B)** *S. aureus* (AH133-GFP) biofilms in LB, FBS and IVWM. To reduce the role of surface attachment, and better explore the role of the different media conditions, biofilms were grown in tissue-culture treated microtiter plates.

When grown in LB media, *P. aeruginosa* displayed minimal biofilm formation after 24 hours (Figure 4B). This is possibly due to the tissue culture treatment of the wells, which is known to reduce the surface attachment of bacteria. On the other hand, in FBS and IVWM, 24-hour old *P. aeruginosa_biofilms* were observed as mat-like structures; the average thickness was 41 ±2 μm in FBS, and 39±1 μm in the IVWM (Figure 4B). Notably, this highlights the role of host components (as in FBS and in the IVWM) in *P. aeruginosa* biofilm formation.

Despite the fact that *S. aureus* biofilms are thinner in the IVWM as compared to that in FBS (Figure 4A), the MBEC_80_ in the IVWM is higher (>512 μg/mL) than that measured in FBS (16 μg/mL) (Table 2). Previous studies have correlated the thickness of biofilms to the antibiotic resistance observed, with thicker biofilms typically observed to display increased antibiotic tolerance, possibly due to reduced antibiotic penetration and the production of deeper gradients [114,115]. However, the presence of serum along with additional host components such as collagen, has been observed to increase antibiotic tolerance for *P. aeruginosa* and *S. aureus* aggregates [23,115–119].

### 3D Biofilm structure of *P. aeruginosa* and *S. aureus* co-species biofilms

To examine co-species biofilms, *P. aeruginosa* (PAO1-mCherry) and *S. aureus* (AH133-GFP) were inoculated in a 1:1 ratio, followed by *in situ* visualization of 24-hour biofilms (Figure 5). In LB media, under co-species conditions, *P. aeruginosa* displayed minimal biofilm formation (similar to that observed in single-species state) (Figure 5A and B). Notably, under co-species conditions, *S. aureus* was also seen to form sparse biofilms. Based on known interspecies interactions between the two pathogens, it is likely that under co-species conditions in LB media, inoculated *P. aeruginosa* results in killing of *S. aureus*, and therefore significantly reduced *S. aureus* biofilms are observed [85,120].

**Figure 5:**
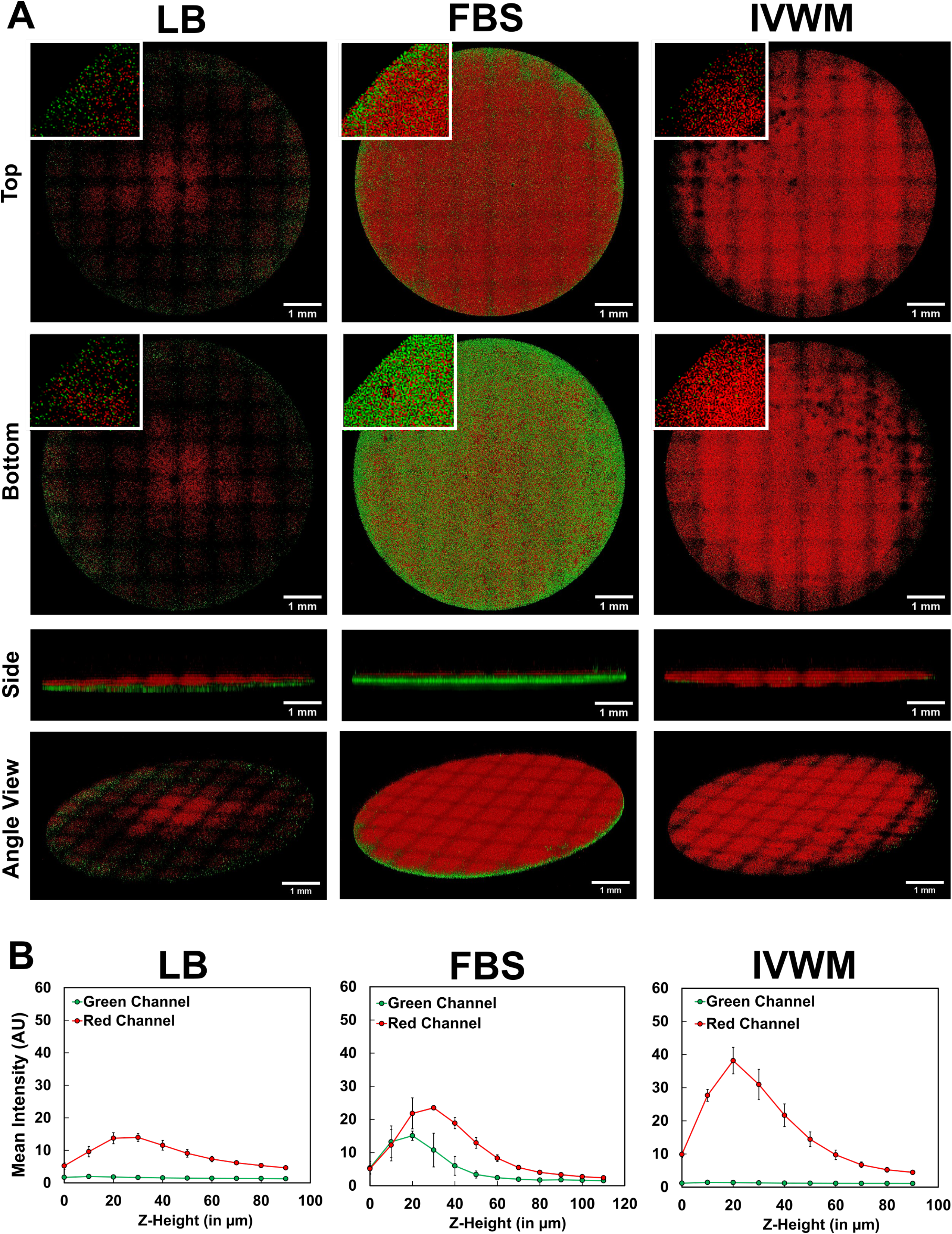
3D biofilm structure of co-species *P. aeruginosa* and *S. aureus* biofilms in IVWM shows distinct predominance of *P. aeruginosa*. **(A)** Tile-scan confocal microscopy of *P. aeruginosa* (PAO1-mCherry) and *S. aureus* (AH133-GFP) co-species biofilms in LB, FBS and IVWM. **(B)** Mean intensity of fluorescence (representing PAO1-mCherry and SA-GFP) across the Z-height of the biofilm (with the bottom as Z = 0). Error bars represent SEM, n=3.

In FBS, the two pathogens were observed to form a robust co-species biofilm (Figure 5A and B), consisting of a thick mat of *S. aureus* (thickness 20±2 μm) and *P. aeruginosa* (thickness 27±1 μm). This indicates that presence of host components (as in FBS), enables the coexistence of *S. aureus* and *P. aeruginosa* biofilms, which is in accordance with previous work that reports the ‘rescue’ of *S. aureus* biofilms [81], and protection from killing by *P. aeruginosa*, in the presence of host components.

However, this is notably different in the presence of additional host components, that more closely mimic the wound milieu. In the IVWM, the co-species biofilm shows a distinct predominance of *P. aeruginosa* (average thickness 29±3 μm), with the presence of *S. aureus* significantly less as compared to that observed in FBS (Figure 5A and B). This could result from the presence of additional host factors countering the protective effect against *P. aeruginosa* killing, or that certain factors in the IVWM, such as lactoferrin exhibit antimicrobial activity [53,121,122], including against *S. aureus* biofilms [51,112,123–125].

Overall, our results indicate that the IVWM supports the formation of dense, mat-like biofilms of *P. aeruginosa* and *S. aureus* when grown alone, and under co-species conditions the polymicrobial biofilm shows a distinct predominance of *P. aeruginosa*. This is similar to *in vivo* conditions where in spite of well-established co-existence of the two pathogens, *P. aeruginosa* appears to outcompete *S. aureus* [18,19].

## Conclusions

Based on previous reports of clinical wound fluid composition, we have developed an *in vitro* wound milieu (IVWM), consisting of fetal bovine serum, with additional host matrix and biochemical factors. Our results indicate that the IVWM recapitulates key *in vivo* biofilm features such as biomass formation, metabolic activity, antibiotic tolerance, three-dimensional structure, and interspecies interactions. Notably, under both planktonic and biofilm conditions, the IVWM supported a distinct predominance of *P. aeruginosa*. This is important to explore further, particularly in clinical and *in vivo* conditions, given that *P. aeruginosa-S. aureus* interactions have been largely studied in *in vitro* systems [20,21,120,123]. Notably, we find that this is distinct from that observed in serum alone, underscoring the role of additional matrix and biochemical factors.

While the IVWM recapitulates key factors in the wound microenvironment, it is certainly not representative of all aspects of the wound milieu. However, its ease of formulation, use of widely-available components, and compatibility with standard biofilm assays, lends itself well for further adaptations and modifications, such as the inclusion of additional factors such as commensal microbes, glucose, matrix-metalloproteinases [126–130]. Given this, the IVWM holds potential as a tractable approach to study wound biofilms under host-relevant conditions, particularly for high-throughput applications such as time-lapse biofilm studies and combination antimicrobial approaches. In doing so, it could bridge the gap between reductionist *in vitro* systems and *in vivo* models, and provide more human-relevant insights in laboratory biofilm studies.

## Acknowledgements

We thank Drs. Derek Fleming and Kendra Rumbaugh (Texas Tech University Health Science Center, Lubbock) for the bacterial strains, Dr. Avinash Sharma (National Center for Microbial Research, NCCS, Pune) for erythromycin, and Dr. Amit Agarwal for 96-well black plates. We acknowledge Sujaya Ingle, NCL-Innovation Park, for technical assistance with confocal microscopy and Dr. Kaspar Kragh for inputs on the tile-scan approach. We also thank Drs. Ameeta Ravikumar, Shadab Ahmed, Tuli Dey and Geetanjali Tomar (Institute of Bioinformatics and Biotechnology, SPPU) for use of certain laboratory equipment.

## Author Contributions

**Snehal Kadam:** Conceptualization, Methodology, Investigation, Validation, Formal analysis, Data curation, Visualization, Writing original draft, Editing draft. **M Vandana:** Conceptualization, Methodology, Investigation, Validation. **Devyani Bhide:** Investigation, Validation. **Rutuja Ugale:** Investigation. **Utkarsha Tikhole:** Investigation. **Karishma S Kaushik:** Conceptualization, Methodology, Formal analysis, Project administration, Supervision, Writing original draft, Editing draft, Funding acquisition.

